# Coordination of movement via complementary interactions of leaders and followers in termite mating pairs

**DOI:** 10.1101/2021.03.05.434098

**Authors:** Nobuaki Mizumoto, Sang-Bin Lee, Gabriele Valentini, Thomas Chouvenc, Stephen C. Pratt

## Abstract

Leadership of animal group movements depends on social feedback, hence leader’s signals and follower’s responses should be attuned to each other. However, leader and follower roles are difficult to disentangle in species with high levels of coordination. To overcome this challenge, we investigated a simple case of movement coordination: termite pairs in which a female leads a male as they search for a nest site. To tease apart leader and follower roles, we created conspecific and heterospecific pairs of *Coptotermes gestroi* and *C. formosanus*, which share a pairing pheromone so that males follow females of either species. Conspecific pairs were stable for both species, even though *C. gestroi* females produce less pheromone than *C. formosanus*. Heterospecific pairs with *C. gestroi* males were also stable, but not those with *C. formosanus* males. We attributed this difference to the *C. gestroi* male’s unique capacity to follow females that release small amounts of pheromone; *C. formosanus* males cannot follow or reject *C. gestroi* females as unsuitable. This conclusion was supported by an information-theoretic analysis that detected information flow from *C. formosanus* females to *C. gestroi* males as in conspecific pairs, but not from *C. gestroi* females to *C. formosanus* males. Despite their following ability, *C. gestroi* males lost to *C. formosanus* males in competitions to follow *C. formosanus* females. Thus, partner selection has shaped the species-specific association of mating pairs. Our results demonstrate that a similar level of coordination can emerge from distinct sets of complementary sender-receiver interactions.

## Introduction

Animals often move as a group while searching for a safe place or feeding site. Coordinated group movements are achieved by rules for interactions among group members, with individuals often playing different roles [1,2]. One or a few individuals initiate movement, and other members follow the leader [3,4]. Such leadership strongly affects the collective outcome of group movements [5]. When a pair of individuals explore the environment together, a leader-follower relationship is almost inevitable; the first to move is the leader, and the other has no option but to follow [6]. Thus, many studies on pairs have focused on how partners respond to each other to control movement speed and turning angle [7–10]. As successful coordination results from social feedback, innate behavioral differences between partners can promote or hinder coordination [11]. Especially if the pair shares a common goal, leader phenotypes should complement follower phenotypes to maintain stable coordination, resulting in a species-specific manner of social interaction.

Tandem running in termites is among the simplest leader-follower relationships. Unlike ants, where tandem runs recruit colony members to specific resource locations [12,13], termite mating pairs perform tandems after dispersal, while seeking sites for colony foundation [14]. The female leads the tandem and releases a short-range sex pheromone to guide the male [15,16], and the male touches the female’s abdomen with its antennae and mouthparts, indicating its continued presence [14,17]. As the sex pheromone varies among species [15], female behavior can also vary in order to transmit species-specific signals efficiently [18]. We predict that males coevolved species-specific following capacity to form stable tandems with conspecific females.

To disentangle the contributions of leaders and followers to behavioral coordination, we made heterospecific pairings between related species. *Coptotermes gestroi* (Wasmann) and *C. formosanus* (Shiraki) evolved in allopatry, but both are now invasive and found in sympatry in some coastal cities [19–21]. In south Florida, USA, heterospecific mating events have been observed, resulting in hybrid colonies [22]. Heterospecific pairing can occur because the species share the same pairing (sex) pheromone, (3Z,6Z,8E)-dodeca-3,6,8-trien-1-ol, emitted from tergal glands at the tip of the abdomen of females [23] (Fig. 1A). The main difference is the quantity of pheromone, and thus the strength of the transmitted signal; *C. formosanus* females produce ~10x more pheromone than *C. gestroi* females [23]. Based on this difference, we hypothesized that males of these species evolved different tandem following capacities matched to their conspecific female’s signal strength.

**Figure 1.**
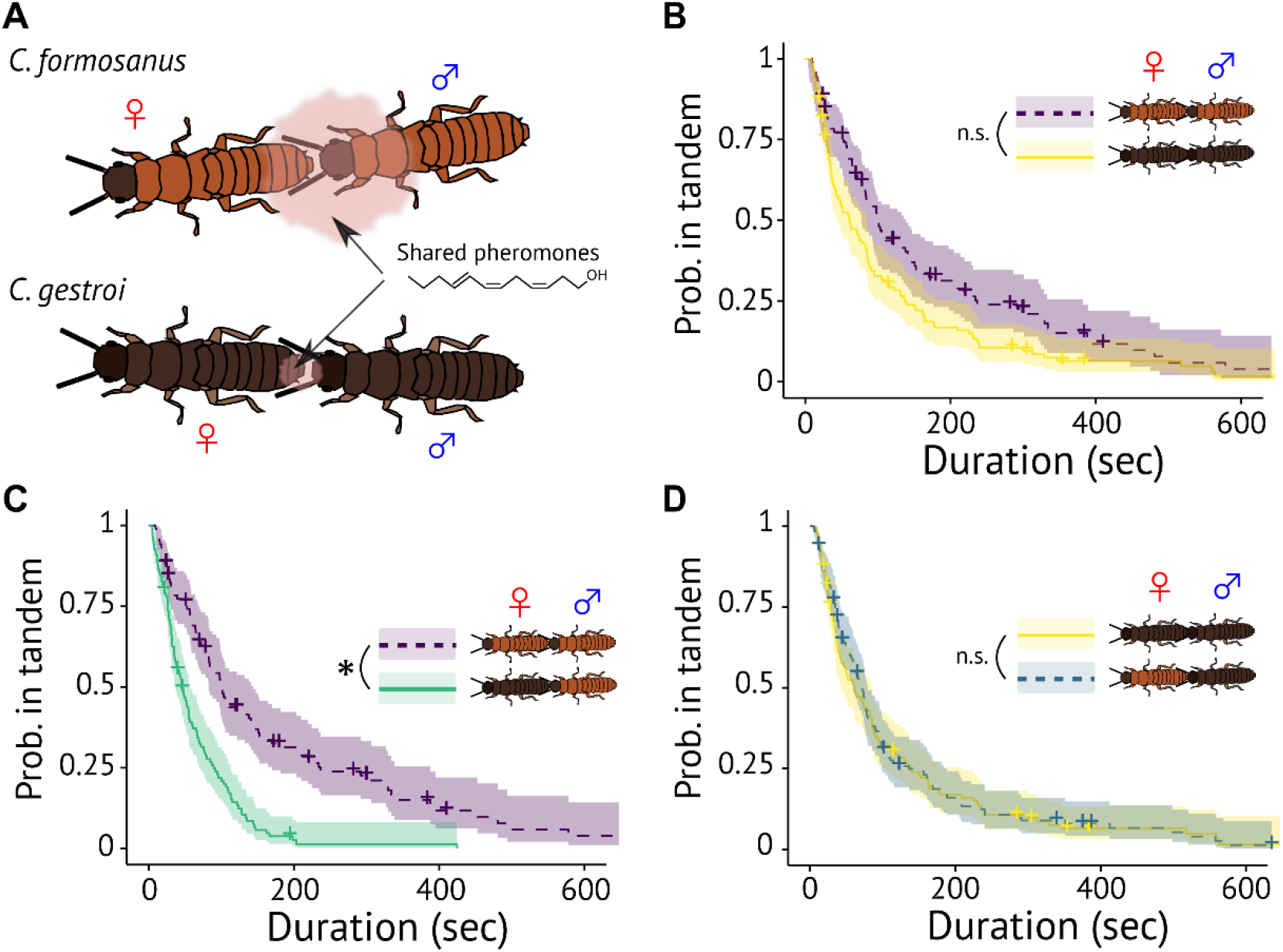
Interspecific variation of tandem running in *Coptotermes* termites. (A) During tandem runs, the female leader releases a short-range pheromone that attracts the male follower. The chemical is shared between *C. formosanus* and *C. gestroi*, but the quantity is much larger in *C. formosanus*. (B-D) Comparison of the duration of tandem running until separation across different combinations. Kaplan-Meier survival curves were generated for each pairing combination. * indicates significant difference (mixed effect Cox model, *P* < 0.05). + indicates censored data due to the end of observations. Shaded regions show 95% confidence intervals. Shaded regions show 95% confidence intervals.

## Results

Termites were collected during synchronized dispersal flights of the two species in south Florida. We observed tandem running behavior of single male/female pairs in a petri-dish arena; all pairing combinations of the two species were observed. Despite the relatively small quantity of sex pheromone involved, we found that *C. gestroi* conspecific tandem runs lasted as long as those of *C. formosanus* (mixed-effects Cox model, χ^2^_1_ = 0.942, *P* = 0.332, Figure 1B). As for heterospecific tandem runs, their durations were asymmetric. When the male was *C. gestroi*, heterospecific tandems lasted as long as conspecific ones (mixed-effects Cox model, χ^2^_1_ = 0.01, *P* = 0.91, Figure 1D). When the male was *C. formosanus*, heterospecific tandems ended sooner than conspecific ones (mixed-effects Cox model, χ^2^_1_ = 19.52, *P* < 0.001, Figure 1C). Thus, tandem runs were unstable only for the combination of a *C. gestroi* female and a *C. formosanus* male, as predicted in [22,23].

Next, to investigate the role of females and males in heterospecific coordination, we observed the moving speed during tandem runs. Moving speeds of females and males were highly correlated across all pairing combinations (Fig. S1). However, modal moving speed was higher for conspecific *C. formosanus* pairs (17.0 mm/sec) than for conspecific *C. gestroi* pairs (14.5 mm/sec, Fig. S2). Thus, for heterospecific pairs to synchronize their movement, one or both partners need to adjust their speed. We found evidence that males make this speed adjustment; the modal speed of heterospecific pairs was similar to that of the female’s conspecific tandem runs (*C. formosanus* female-*C. gestroi* male: 17.5mm/sec, *C. gestroi* female-*C. formosanus* male: 14.5mm/sec, Fig. S2). Also, across all tandem runs, speed depended on the female species (LMM; female moving speed, female species: χ^2^_1_ = 14.888, *P* < 0.001, male species: χ^2^_1_ = 2.0802, *P* = 0.1492; male moving speed, female species: χ^2^_1_ = 12.2442, *P* < 0.001, male species: χ^2^_1_ = 1.5145, *P* = 0.21845).

The asymmetry between heterospecific pairings was further supported by an information-theoretic analysis. We used transfer entropy to quantify the degree to which the female leader’s motion predicts that of the male follower, a measure of coupling strength within the pair. Transfer entropy quantifies how well knowledge of present behavior of the sender reduces uncertainty about the future of behavior of the receiver, after taking account of the receiver’s history [24,25]. This value can be determined for both directions, with the difference giving a measure of the net direction and amount of information flow. An earlier study on conspecific tandem runs by *C. formosanus* showed that the net flow was from leader to follower; that is, the female largely determines both the direction and speed of the male [9]. This pattern is consistent with the female carrying out a random search of the environment and the male simply following closely behind her. Thus, successful heterospecific tandems would show a similar pattern, but that unstable tandems would show low net transfer entropy from female to male.

We calculated transfer entropy after coarse-graining the spatial trajectories of termites into sequences of discrete movements (pause, clockwise turn, or counterclockwise turn). As expected, the future behavior of males was significantly predicted by the present behavior of females in all stable combinations—that is, conspecific pairs or heterospecific pairs of *C. formosanus* females and *C. gestroi* males (Wilcoxon rank-sum test, *P* < 0.001) (Fig. 2, Table S2). In these pairs, information flow from females to males was significantly stronger than in the opposite direction (Wilcoxon signed-rank test, *P* < 0.05) (Table S2). However, in the unstable heterospecific pairs of *C. gestroi* females and *C. formosanus* males, neither female nor male behavior was significantly predicted by their partner’s behavior (Wilcoxon rank-sum test, *P* > 0.05); thus, there was no predominant direction of information flow (Wilcoxon signed-rank test, *P* > 0.05) (Fig. 2, Table S2). The lack of predictive power by female behavior of male behavior indicates deficient following by males in this combination.

**Figure 2.**
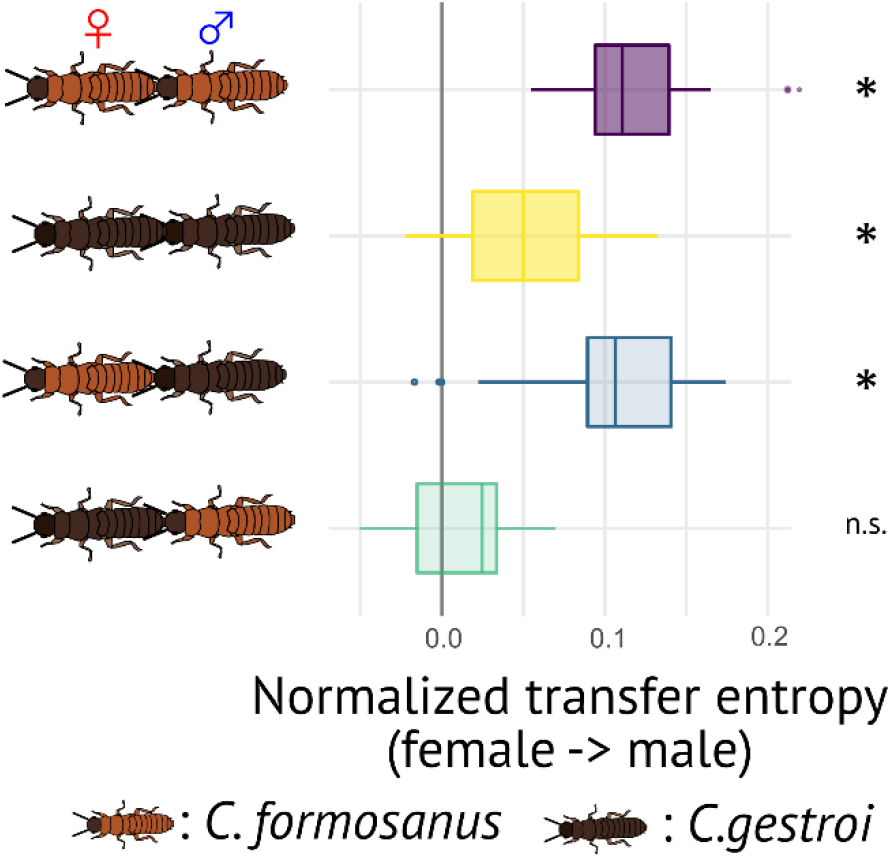
Comparison of the strength of information flow during tandem runs. The predominant direction of predictive information is given by the proportion of uncertainty reduction explained by the interaction between leading females and following males. * indicates the combination with significant information flow from female to male.

These results suggest that *C. formosanus* males do not maintain heterospecific tandem runs because they only follow females that release large amounts of sex pheromone. In contrast, *C. gestroi* males are accustomed to small quantities and are not challenged in following the larger amount released by *C. formosanus* females. Males of *C. formosanus* may have difficulty detecting small amounts of pheromone, or they may instead lack the motivation to follow weak pheromone signals. To investigate their motivation, we observed how males behave when they become separated from their leader. After separation, the female pauses while the male engages in active local search, and this dimorphism enhances re-encounter rates [18]. By moving slowly just after separation, males increase their re-encounter rate with the same partner. By instead moving quickly, they can more efficiently search for a new partner [26]. Thus the male’s movement speed right after separation can indicate a male’s evaluation of the female (much as waiting time of separated leaders reflects follower quality in ant tandem runs [26]). Slow movement indicates a relatively positive evaluation of the separated female and preference to re-unite; fast movement means a lower rating and a preference for finding a new partner.

After separation, females of both species slowed down significantly irrespective of partner species (comparison of mean speed two seconds before and two seconds after separation, LMM, *P* < 0.01, Fig. 3ABDE). Males of *C. gestroi* evaluated both conspecific and *C. formosanus* females as good leaders because they slowed down just after the separation to enhance re-encounter rates (LMM, *P* < 0.05, Fig. 3DE). On the other hand, *C. formosanus* males slowed down upon separation from conspecific females (LMM, estimate±s.e. = −0.4409±0.2217, χ^2^_1_ = 3.9546, *P* = 0.04674, Fig. 3A), whereas they increased their speed after separating from *C. gestroi* females (LMM, estimate±s.e. = 1.1553±0.1981, χ^2^_1_ = 34.003, *P* < 0.0001, Fig. 3B). Moreover, when the original partners did re-encounter each other, their probability of resuming a tandem run was lower for pairings of a *C. formosanus* male and a *C. gestroi* female than for other pairing combinations (GLMM, Tukey’s test, *P* < 0.01, Fig. 3CF). These results suggest that a *C. formosanus* male evaluates a *C. gestroi* female as a poor leader and begins to search for another partner upon separation. Thus, male preference plays an important role in the success of heterospecific tandem runs.

**Figure 3.**
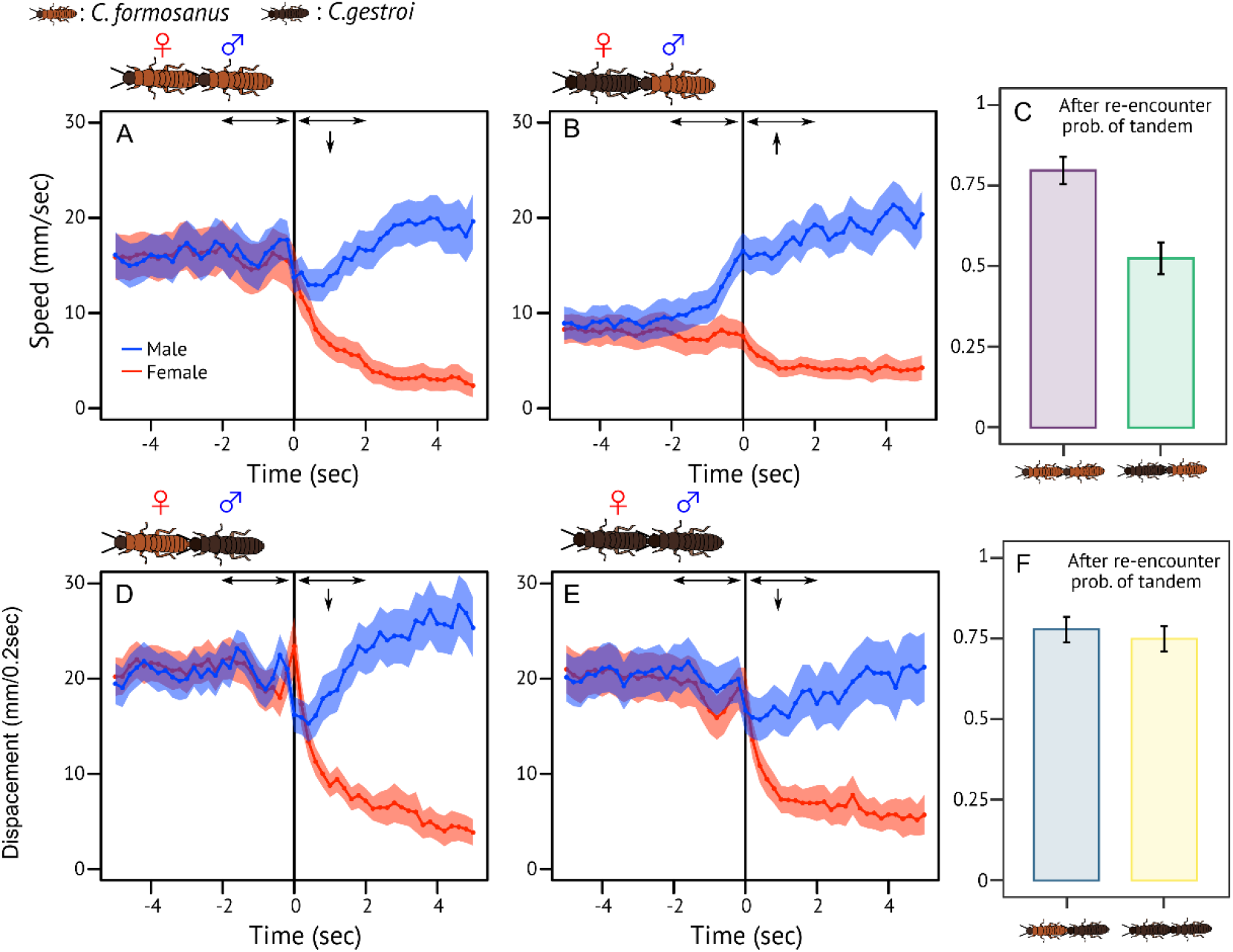
Behavioral change of tandem runners before and after separation events. (A,B,D,E) The time development of speed across different combinations of mating pairs. Pair separation occurs at 0 sec. Shaded regions indicate mean speed ± 2*SE*. Arrows indicate significant increase (upwards) or decrease (downwards) of male speed compared with before separation. (C, F) The probability of resumption of a tandem run upon re-encounter following a separation event. Bars indicate the mean ± *SE*.

In natural conditions, synchronized swarming events often result in a high density of mate searchers, where multiple males attempt to form a tandem with a single female [27]. Because tandem runs involving three individuals are unstable, males compete for the position behind a female [28]. For these two species, heterospecific competition is predictable as *C. formosanus* females are attractive to males of both species. Thus, we investigated the outcome when one *C. formosanus* male and one *C. gestroi* male compete over one *C. formosanus* female.

In these cases, the termites could be in one of four different states (Fig. 4A): search (no tandem run), conspecific tandem run of *C. formosanus*, heterospecific tandem run of *C. formosanus* female and *C. gestroi* male, and three-partner tandem run with the two competing males side by side behind the female. When two individuals were in tandem, there was no interspecific difference in the probability to return to the search state (Fisher’s exact test, *P* = 0.5174). However, when three individuals were in tandem, the probability of state transition was asymmetrical; competitions were twice as likely to end with victory by the *C. formosanus* male than by the *C. gestroi* male (Binomial test, *P* < 0.001). As a result, the most frequently observed state was conspecific tandem running by *C. formosanus* (Fig. 4B). These results demonstrate that *C. formosanus* males are superior to *C. gestroi* when competing over a female, possibly because of their better-matched moving speed and body size.

**Figure 4.**
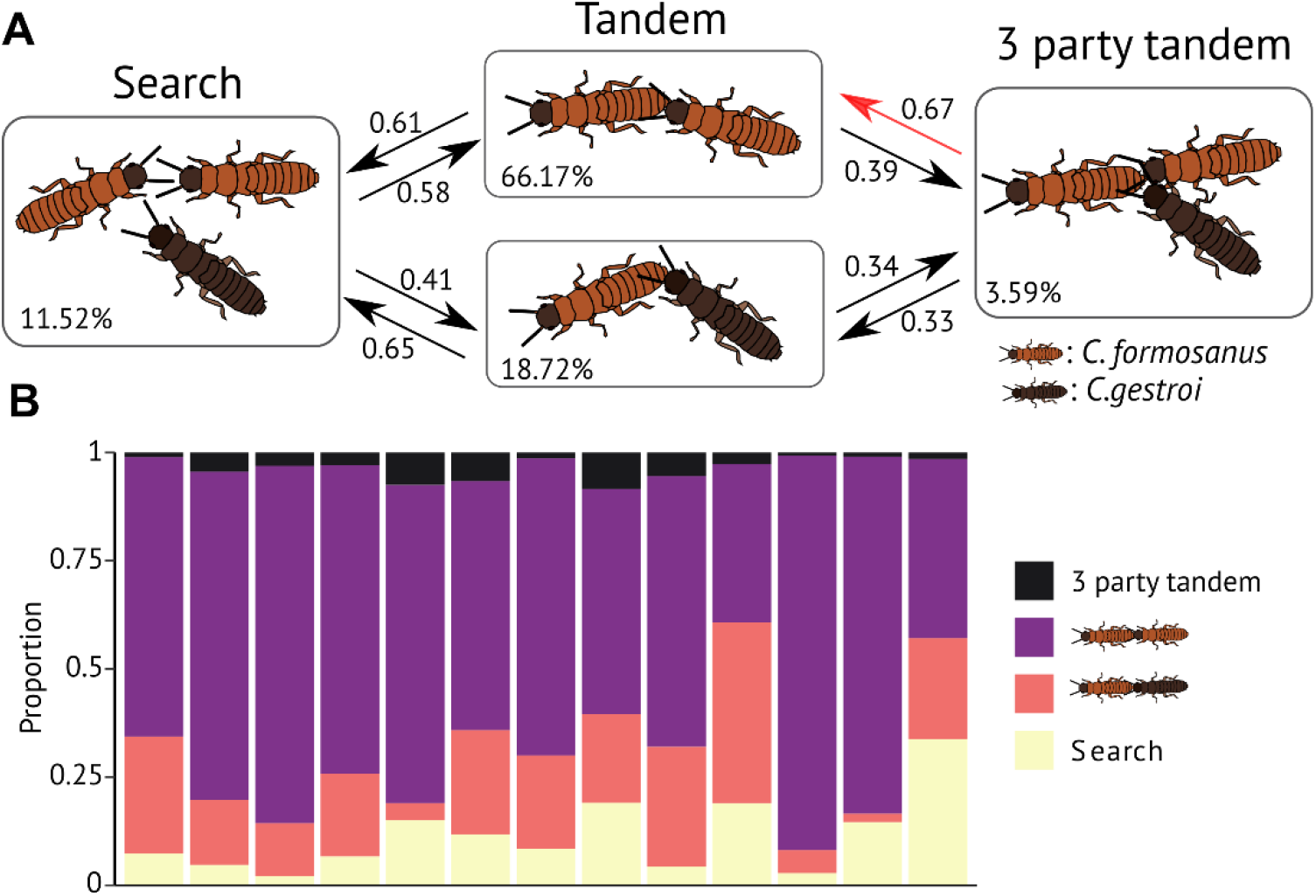
Competition between *C. formosanus* and *C. gestroi* males to follow one *C. formosanus* female. (A) State transition diagram for a 3-party tandem run with a *C. formosanus* female, a *C. formosanus* male, and a *C. gestroi* male. The transition from search to 3-party tandem is not shown (~ 0.01). The percentage in each box indicates overall time in that state. (B) Proportion of time in each state during observation. Each bar indicates replicate. Overall, the conspecific tandem of *C. formosanus* was the most frequent state.

## Discussion

Our study has demonstrated that tandem coordination depends on a close association between the behavior of male followers and the signal strength of female leaders. Males of *C. gestroi*, whose females produce only a tenth of the amount of sex pheromone as *C. formosanus* [23], are adapted to follow a weak signal and so can maintain stable tandem runs with *C. formosanus* females. On the other hand, *C. formosanus* males are adapted to follow females with stronger signals and display a poor ability to maintain tandem runs with *C. gestroi* females. When they become separated from a female with a weak signal, *C. formosanus* males search for an alternative leader. This result demonstrates active behavioral mate choice in termites, hypothesized in previous studies but not shown empirically [29,30]. Note that females behaved the same regardless of male species, implying that they maintain the tandem passively through pheromone production, rather than actively choosing their potential mate. In summary, our results suggest that behavioral coordination in termite tandem runs is a product of coevolution between females and males. The species-specific association of leader and follower phenotypes may explain previous observations on the collective behavior of mixed-species groups; some function as well as conspecific groups, while others show a loss of coordination [31–33].

Leadership may be more likely in some individuals, due to traits like body size or personality (reviewed in, e.g., [6,34,35]). In such a group, coordination may be difficult when there is a conflict of interest among members (e.g., about where to go). However, this is not the case in termite tandem runs. There is no conflict over leadership because roles are usually fixed by sex, with females leading and males following, although this is flexible in occasional same-sex pairs [28]. Additionally, it is reasonable to assume that the pair share the same goal of successful colony foundation. Predation risk is high during tandem [36], so pairs must establish a nest as soon as possible [37]. Their highest priority is not who they found a colony with but simply to found as soon as possible [38]. Thus, we conclude that unstable tandem runs result not from conflicts of interest but from a communication mismatch between *C. gestroi* females and *C. formosanus* males, where female signaling is not sufficient for the males to follow.

Laboratory experiments have shown that individual behavior underlying group coordination can evolve in just a few generations [39,40]; however, such changes have yet to be documented in the field. Species invasions provide opportunities to observe evolutionary changes in behavior [41,42]. In our study of an invasive population, we found less stable tandem runs than those previously observed in a native population of *C. formosanus* in Japan ([18], Supplemental text, Fig. S4). This suggests modification of tandem coordination following invasion, although such differences may have already existed between the source population and other native ranges [43]. Further investigation is needed to determine if properties of the invasive population (e.g., high density or relatively low genetic diversity [43]) are causes of behavioral change in *C. formosanus* from Florida. Moreover, colony foundation experiments have confirmed that hybrid colonies can last > 2 years [44], raising questions about potential behavioral changes of hybrid adults.

Evolutionary theory predicts that the development of conserved morphological structures can involve significant modifications in their regulatory mechanisms [45,46]. We argue that this is possible even in behavioral systems. For example, similar group-level patterns can emerge from different individual-level behavioral rules regulating social interactions [47,48]. In this study, we show that a similar level of behavioral coordination can be achieved from different leader/follower combinations. In *C. formosanus,* the leader produces an abundant signal tracked by a competitive follower; while in *C. gestroi,* the leader produces a weaker signal, but the follower has enhanced tracking ability. Tandem runs are seen across most termite taxonomic groups [15]. However, our results imply that their underlying mechanisms for coordination can vary, because the adaptiveness of a tandem run is not determined by how they coordinate but by how well they maintain contact during nest site search in a vulnerable period. By emphasizing that there are multiple solutions for the same coordination problem, our study has implications beyond pair coordination and gives insight into the convergent evolution of collective behavior across different taxa.

## Materials and Methods

### Termites and experimental arena

Two *Coptotermes* termites, *C. formosanus* and *C. gestroi*, evolved in allopatry in the course of 18 million years of evolution [49]. However, due to increasing global human activity, they have spread and now occur in sympatry in three distinct geographical regions: Taiwan, Hainan, and Southeast Florida [22]. We collected alates of *C. formosanus* and *C. gestroi* using a light-trapping system at dusk between Apr 18th and 20th in 2020 in Broward County (Florida, USA). We brought the alates to the laboratory and maintained them on wet cardboard at 28°C. We used individuals who shed their wings by themselves and observed their behavior within 12 hours after the flight. Each individual was used only once.

We performed all observations in an experimental arena made by filling a petri dish (ø=140mm) with moistened plaster. The petri dish had a clear lid during observations. A video camera above the arena was adjusted so that the arena filled the camera frame. We extracted the coordinates of termite movements from all obtained video, using the video-tracking system UMATracker [50]. All data analyses were performed using R v4.0.1 [51].

### Comparing tandem run stability across different pair combinations

To explore interspecies differences in tandem running behavior, we introduced one female and one male to the experimental arena and recorded their behavior for 30 minutes. We tested four different species combinations: conspecific pairs of *C. formosanus* (Cf-Cf), conspecific pairs of *C. gestroi* (Cg-Cg), heterospecific pairs of female *C. gestroi* and male *C. formosanus* (Cg-Cf), and heterospecific pairs of female *C. formosanus* and male *C. gestroi* (Cf-Cg). We prepared ten replicates for each combination.

During observations, termite pairs were in one of three states: (i) tandem running, (ii) interacting but not tandem running, and (iii) searching (two are in a distance). Following a previous study [27], we classified the pairing states based on the coordination of female and male. We defined them as interacting (or tandem running) when the distance between their centroids was less than 1.3 × mean body length. This value was 11.57 for *C. formosanus*, 9.75 for *C. gestroi*, and 10.65 for heterospecific pairs, respectively. We selected this distance to slightly exceed the average body length because termites in a tandem run are nearly in physical contact [18]. An interacting pair was considered to be performing a tandem run only if they met the following criteria [27]. First, the interaction needed to last for more than 5 seconds; a very short separation (< 2 seconds) was not regarded as a separation event unless the distance between individuals was greater than 20 mm. Second, both termites needed to move more than 30 mm while interacting. After separation, we considered that individuals engage in separation search until they interact with an individual again for more than 1 second. We down-sampled all videos to a rate of five frames per second (= every 0.2002 sec) for this analysis.

We obtained 103, 110, 120, and 132 tandem run events for Cf-Cf, Cg-Cg, Cg-Cf, and Cf-Cg, respectively. We compared tandem duration between the two conspecific pairs and between heterospecific and conspecific pairs for each male species. We used the mixed-effects Cox model (coxme() function in the coxme package in R [52]), with female species as a fixed effect and video id as a random effect. The likelihood ratio test was used to determine the statistical significance of each explanatory variable (type II test). Observations interrupted by the end of the video were treated as censored data.

### Moving speed during tandem runs

We compared moving speed during tandem runs across different pair combinations to further explore the role of females and males for behavioral coordination. We first calculated the moving step length between two successive frames at 5FPS. The step length distribution was bimodal, with two peaks around 0 and 3mm (Fig. S2). The two peaks can be regarded as representing pauses and moves, respectively. Based on the histogram of each pairing combination using 0.1mm bins, we obtained the value representing the second peak of moving speed (Fig. S2). Then, we defined thresholds to distinguish movements from pauses by multiplying the value of the 2nd peak by a factor of 0.2 (Cf-Cf: 3.4mm, Cg-Cg 2.9mm, Cf-Cg: 3.5mm, and Cg-Cf: 2.9mm) [18]. A pause was defined as a step length shorter than or equal to the threshold. By removing data for pause durations, we obtained a dataset only including moving speed. Finally, we used a linear mixed model to analyze moving speed, where the species of female and male were included as fixed effects and video id as a random effect. The likelihood ratio test was used to determine the statistical significance of each explanatory variable (type II test). Note that, although we present results applying species-specific thresholds, we reached the same conclusions when we used one identical threshold (=2.9mm, obtained from the histogram of a pooled dataset).

### Information transfer between females and males

We examined information transfer between tandem pair members by coarse-graining their movement trajectories into a sequence of discrete behaviors. During tandem runs, the female explores the environment to look for a potential nest site with the following male [14]. In a random search, both move/pause patterns and turning patterns link to search efficiency [53]. We discretized trajectories of each runner to obtain time-series describing the pausing and rotation pattern [9]. The behavior of each runner was classified into three states: pause (P), motion with clockwise rotation (M-CW), and motion with counterclockwise rotation (M-CCW). The pause state was distinguished from others using the threshold obtained in the moving speed analysis. As this threshold was computed on the basis of data sampled at 5FPS (sampling period = 0.2002s), we simply rescaled this threshold by the ratio of sampling periods to obtain that for other sampling periods. If the step length between successive frames was shorter than the threshold, the state of the frame was recorded as a pause P. Otherwise, the state was either M-CW or M-CCW depending on the direction of motion computed as the cross product of movement vectors between successive time steps. If no rotation was detected (i.e., cross-product equal to 0), the rotation direction was copied from the previous time step.

We employed transfer entropy to investigate the coupling between female leaders and male followers during tandem runs (refer to Valentini et al. (2020) for a detailed description of this methodology). Transfer entropy is an information-theoretic measure that quantifies the predictive power given by knowledge of the present state of an individual about the future state of a different individual. In other word, it measures causal interactions between a sender and a receiver in terms of Granger causality [54]. If *L* and *F* are behavioral sequences representing the leading female and the following male, then transfer entropy from *L* to *F* is defined as

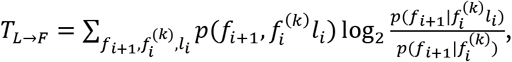

where *l*_*i*_ is the value of sequence *L* at time *i*, *f*_*i*+1_ is the value of sequence *F* at time *i*+1, and 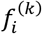 is the *k*-history of *F* at time *i* (i.e., the last *k* states in the sequence). As range of values of transfer entropy is determined by the encoding of states in the sequences, we can normalize its value to obtain a coupling measure in the range of [0;1] by dividing it for its maximum [9,55]. Normalized transfer entropy indicates the proportion of the follower’s future behavior that is predicted by the leader’s present behavior: it is 1 when the follower behavior is entirely determined by the behavior of the leader, and 0 when the two are independent from each other. Furthermore, transfer entropy can be computed in both directions, from leader to follower and from follower to leader. By comparing these values, *T*_*L*→*F*_ and *T*_*F*→*L*_, we can obtain the predominant direction of information flow. The difference in transfer entropy between the two directions, *T*_*L*→*F*_ − *T*_*F*→*L*_, is called net transfer entropy [9,55]. The value is positive when information flow from leader to follower is predominant (*T*_*L*→*F*_ > *T*_*F*→*L*_) and negative when flow from follower to leader (*T*_*L*→*F*_ < *T*_*F*→*L*_) predominates.

To test that our results were the significant, and not an artefact of finite sample size, we artificially created surrogate datasets by pairing time series obtained from leaders and followers not tandem running together; then, we computed transfer entropy for these datasets and compared it with the experimental results [9,55]. To produce a surrogate dataset, we paired randomly selected leaders and followers belonging to different tandem runs. Although females and males from different tandems are still influenced by the same environmental cues of the experimental arena, this randomization process ensures that there are no causal interactions within the surrogate pair. For each pairing combination and parameter configuration, we repeated the randomization process and obtained 100 surrogate datasets. We used these to estimate the mean and standard error of transfer entropy for surrogate datasets with the same size of the experimental ones. Finally, measurements of transfer entropy for the experimental data were discounted by a correction factor given by the mean values estimated from surrogate datasets to account for their finite sample sizes.

Our information-theoretic analysis depended on setting the values of two parameters: the sampling period of continuous spatial trajectories and the history length of transfer entropy, *k*. The optimal choice of these parameters varies for different pairing combinations and focal behavioral patterns due to behavioral, morphological, and cognitive differences manifesting at different time scales [9]. To find good parameterizations, we computed net transfer entropy for 900 different parameter configurations for each species (history length *k* ∈ {1, … , 20} and sampling period {0.0334*s*, … , 1.5015*s*}). The resulting landscapes of net transfer entropy show robustness to different parameter values over most of the tested range (Fig. S3). We selected the parameter configurations that maximize the net transfer of information (Table S1).

For the chosen parameter configurations, we performed two statistical tests. First, we tested if the experimental data showed significantly greater values of transfer entropy with respect to the surrogate data. We used one-sided two-sample Wilcoxon rank-sum tests with continuity correction. Second, we tested differences in the flows of information between the two possible directions (from leaders to followers and from followers to leaders) to determine which among the leader and the follower was the predominant source of information. We used one-sided paired Wilcoxon signed-rank tests with continuity correction. All information-theoretic measures were computed using the rinform-1.0.1 package for R [56].

### Moving speed after separation

When termites in a pair are accidentally separated, females pause while males move to enhance the chances to reunite [18]. As moving speed is related to reunion efficiency [27], we measured the change in movement speed, focusing on a time window around separation events. We compared movement speed between the last 2 seconds before separation and the first 2 seconds after separation. For each separation event, we measured the mean movement speed for both time windows. Then, we used linear mixed models (LMM), with the time window treated as a fixed effect and video ID included as a random effect. The model was fit for each combination of pairs. The likelihood ratio test was used to determine the statistical significance of each explanatory variable (type II test). Finally, we examined if re-encounter after separation resulted in a tandem run or not.

### Interspecific competition over a female

Because males of both species show stable tandem runs with *C. formosanus* females, we introduced one female *C. formosanus*, one male *C. formosanus*, and one male *C. gestroi* to the experimental arena to study interspecific competitions. We prepared 13 replicates and recorded their behavior for 30 minutes. Tandem runs were identified using the method described above. By doing so, we obtained the time series of states observed among three individuals. There were four different states: (i) no tandem run is observed, (ii) tandem run between female *C. formosanus* and male *C. formosanus*, (iii) tandem run between female *C. formosanus* and male *C. gestroi*, (iv) tandem run involving three individuals. We counted tandem runs of three individuals when both males were concurrently interacting with the female. When three individuals were in a straight line, we regarded it as a tandem run of heading female and the male just after her. Then, we counted the number of transitions from one state to another. Usually, state (i) can transit to (ii) or (iii), state (ii) or (iii) to (i) or (iv), state (iv) to (ii) or (iii) (Fig. 4A). Then we compared the tendency of state transition using binomial tests. We also checked if there is a different state transition trend from (ii) or (iii), using Fisher’s exact test.

## Acknowledgments

We thank members of Pratt lab at Arizona State University for helpful discussion, and Michael S. Engel for comments on the manuscript. NM is supported by a JSPS Overseas Research Fellowship and a JSPS Research Fellowships for Young Scientists, SPD and CPD (grant number: 20J00660). GV is supported by research funds from Arizona State University to Prof. Bert Hölldobler. TC is supported by a grant from USDA National Institute of Food and Agriculture Hatch projects number FLA-FLT 005660, and by an NSF-DEB grant, under the agreement No. 1754083.

## Authors contributions

NM: Conceptualization, Methodology, Validation, Formal analysis, Investigation, Data Curation, Writing–original draft preparation, Writing-review and editing, Visualization, Funding acquisition; SBL: Data collection, Methodology, Investigation, Writing-review and editing; GV: Methodology, Validation, Formal analysis, Writing–review and editing; TC: Conceptualization, Methodology, Writing–review and editing, Funding acquisition; SCP: Conceptualization, Writing–review and editing, Supervision

## Supplemental Information

### Supplementary Text

#### Comparison with native population

In a previous study, Mizumoto and Dobata (2019) observed the tandem running behavior of *Coptotermes formosanus* collected in coastal pinewood forest in Wakayama prefecture, Japan [18]. This area is part of the natural geographic range of *C. formosanus*, because there were records of two termitophilous rove beetle species (*Japanophilus hojoi* and *Sinophilus yukoae*), which are specific to *C. formosanus* [57]. The previous observation was for 60 minutes on 17 pairs. To adjust the observation period, we only used the first 30 minutes for the comparison. By applying the same analysis as the present study (Method section), we obtained the duration of each tandem run event. Using a mixed-effects Cox model with the collection area as a fixed effect and the video id as a random effect, we compared the duration of *C. formosanus* tandem runs between native and invasive regions.

We found that tandem runs of pairs collected from the native population lasted much longer than the introduced population (mixed-effects Cox model, χ^2^_1_ = 13.359, *P* < 0.001, Fig. S4). In both regions, mate searchers were collected as they emerged from their nests for swarming for the first time, and they were investigated in our experiments just after the swarming events. Thus, the behavioral difference between native and invasive populations should not be caused by learning or behavioral plasticity. There are several other possible factors. For example, vigor might be reduced in the invasive population because of inbreeding cycles. Indeed, a previous study detected that females with a lower degree of heterozygosity are less likely to be paired with males [29]. If there is a link between pairing pheromone quantity and tandem stability, pheromone quantity released by females might decrease during the invasion process. Also, there may exist differences between native populations, where the strain introduced in Florida might have already shown less stable tandem runs.

**Fig. S1.**
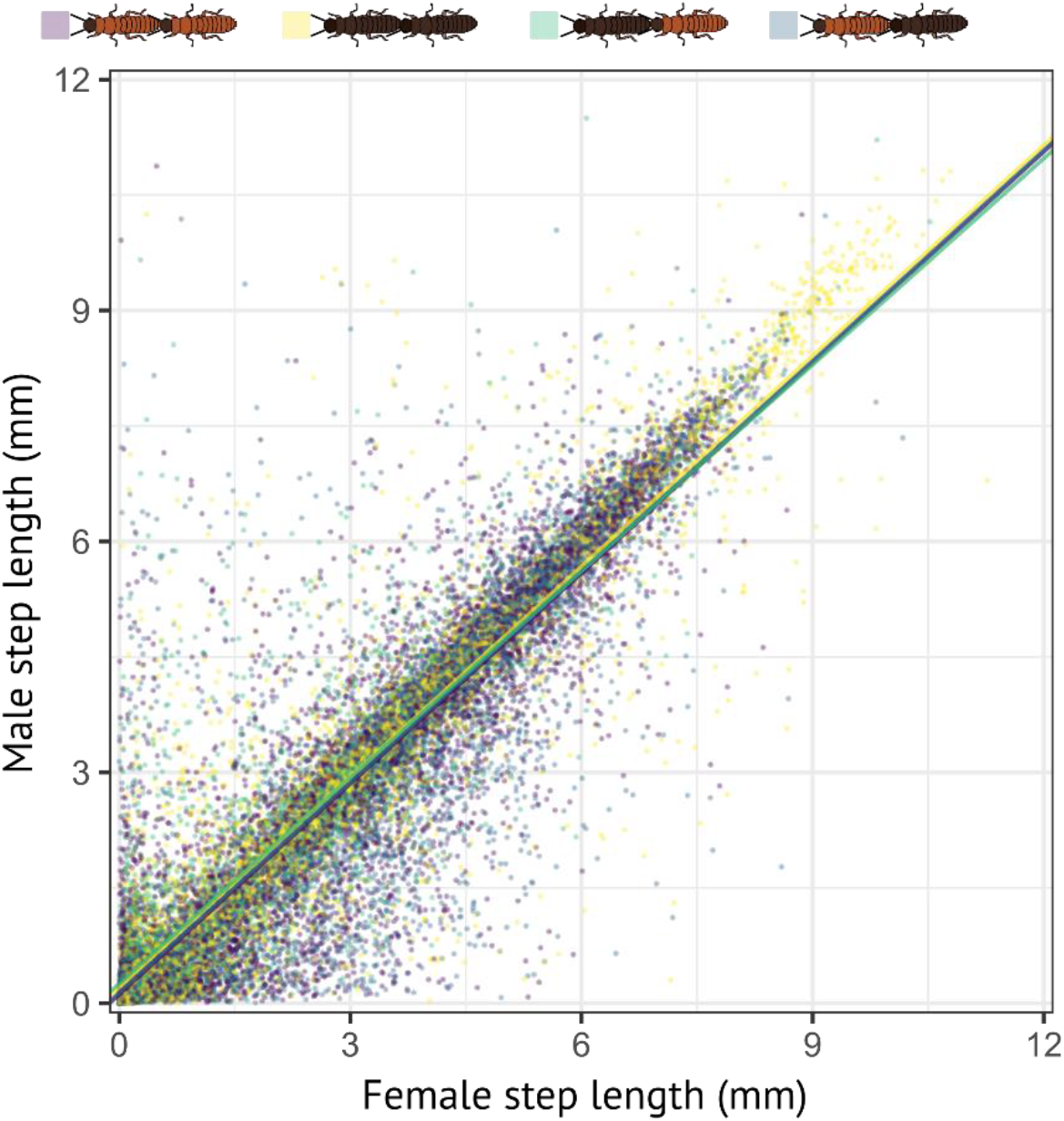
Correlation of step length between females and males during tandem runs. Step lengths were obtained at a rate of 5FPS (every 0.2 seconds). For visualization, we reduced the number of data points by randomly sampling 10% of them.

**Fig. S2.**
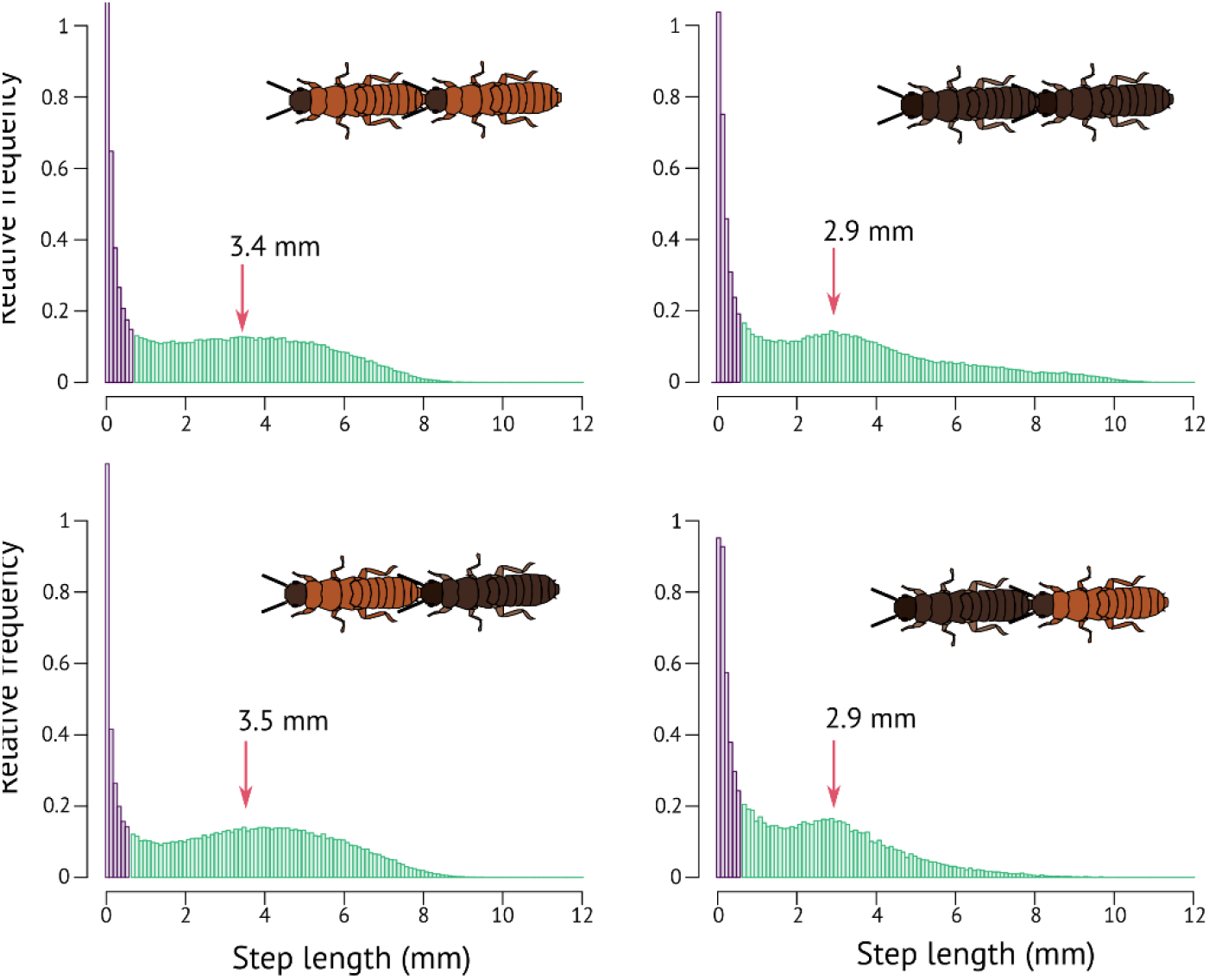
Histogram of the step length between successive frames (0.2s). The histogram was bimodal for all pairs, where the first peak (purple) represents pausing while the second peak (green) represents movement. Red arrows show the peaks of the second histograms.

**Fig. S3.**
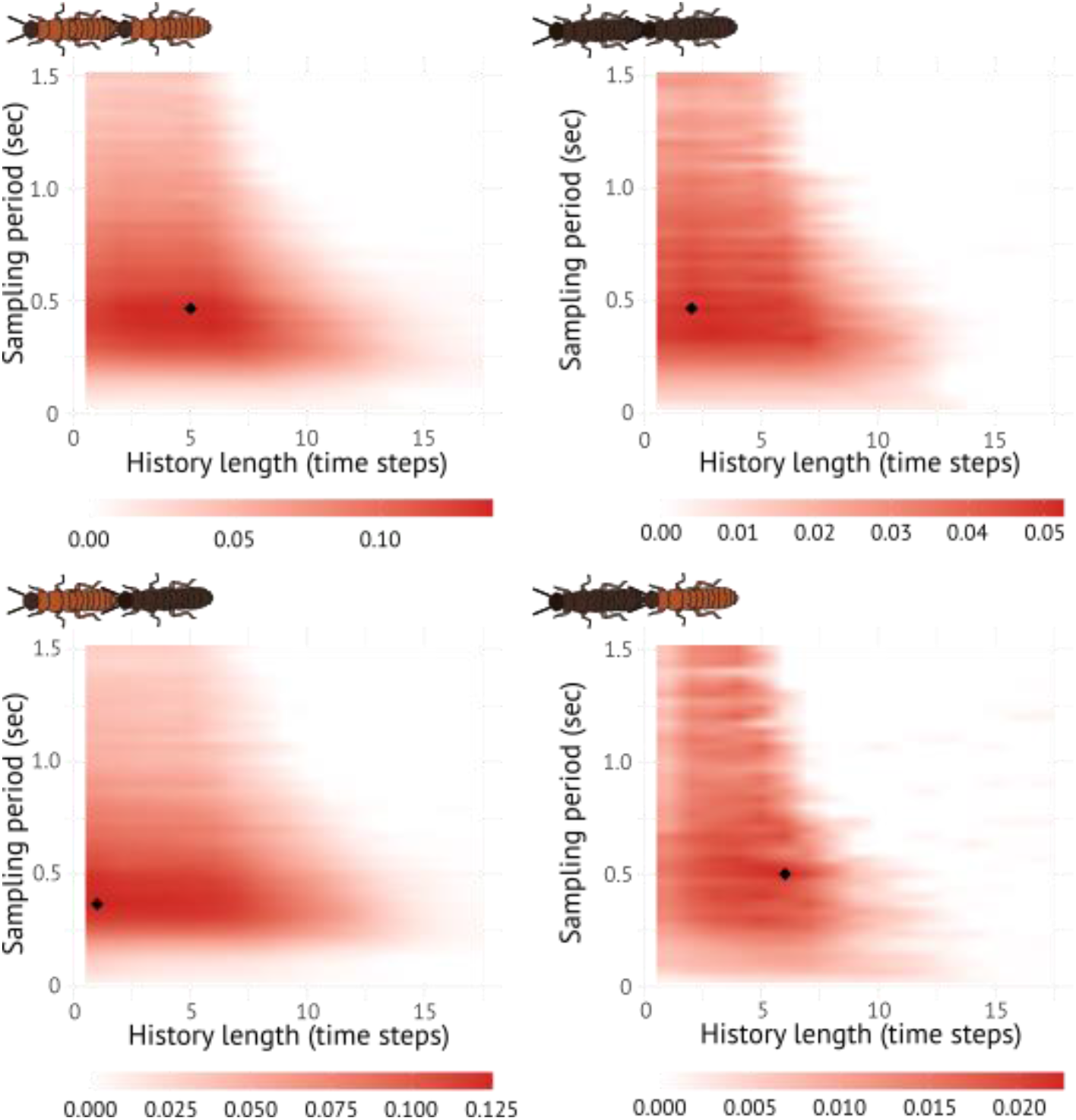
Landscapes of net information transfer. Net transfer entropy (bits) as a function of two parameters: the sampling period and the history length. Colors indicate the intensity of information transfer; the diamond symbol indicates the configuration with maximum magnitude.

**Fig. S4.**
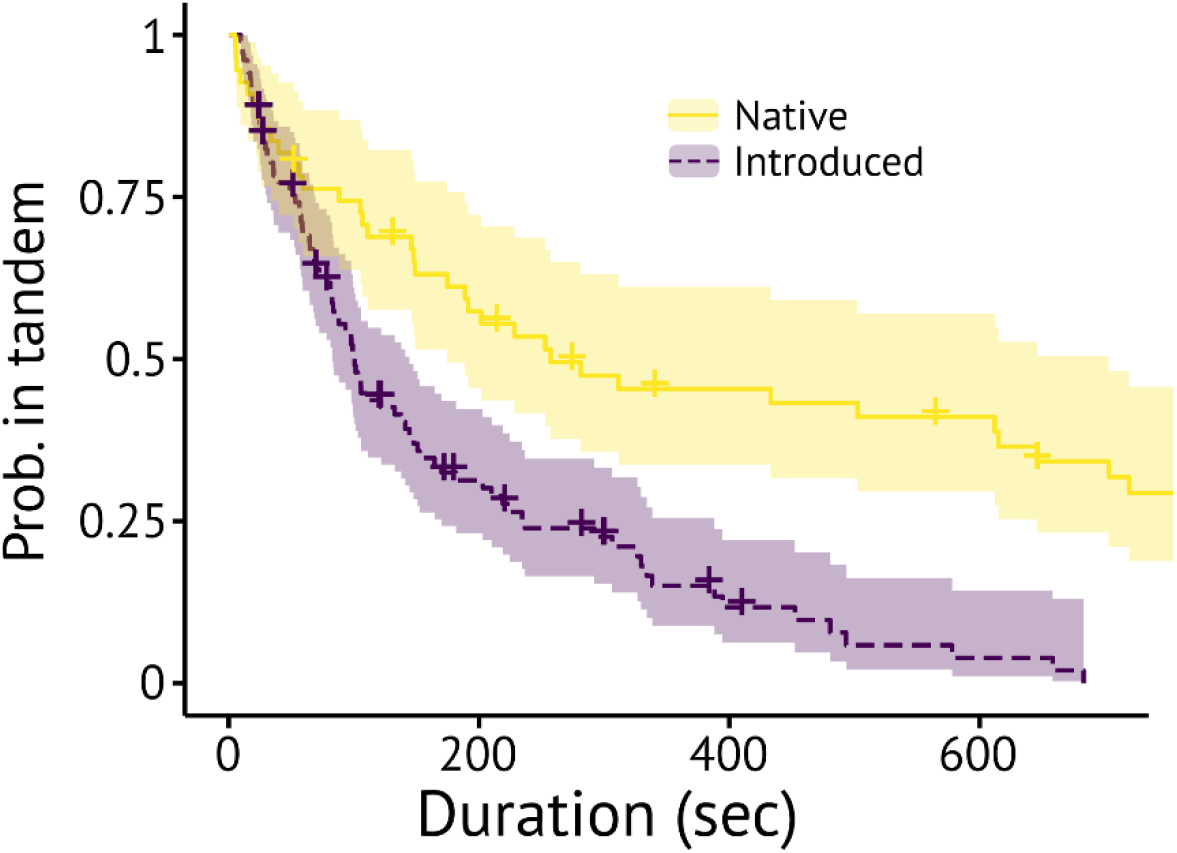
Comparison of tandem stability between the invasive and native population in *C. formosanus*. Kaplan-Meier survival curves were generated for each pairing combination. + indicates censored data due to the end of observations. Shaded regions show 95% confidence intervals. Shaded regions indicate 95% confidence intervals.

**Table S1.** Parameter configurations.

| Female | Male | History length (k) | Sampling period (s) |
| --- | --- | --- | --- |
| *C. formosanus* | *C. formosanus* | 5 | 0.4671 |
| *C. gestroi* | *C. gestroi* | 2 | 0.4671 |
| *C. formosanus* | *C. gestroi* | 1 | 0.3670 |
| *C. gestroi* | *C. formosanus* | 6 | 0.5005 |

**Table S2.** Comparison of information transfer between leaders and followers.

| Female | Male | $T_{L \rightarrow F} > T_{L \rightarrow F}^S$ | $T_{F \rightarrow L} > T_{F \rightarrow L}^S$ | $T_{L \rightarrow F} > T_{F \rightarrow L}$ | $T_{F \rightarrow L} > T_{L \rightarrow F}$ |
| --- | --- | --- | --- | --- | --- |
| *C. formosanus* | *C. formosanus* | $P < 0.001$ ($W = 100$) | $P = 0.109$ ($W = 67$) | $P < 0.001$ ($V = 55$) | $P = 1.0$ ($V = 0$) |
| *C. gestroi* | *C. gestroi* | $P < 0.001$ ($W = 90$) | $P = 0.241$ ($W = 60$) | $P = 0.014$ ($V = 49$) | $P = 0.99$ ($V = 6$) |
| *C. formosanus* | *C. gestroi* | $P < 0.001$ ($W = 90$) | $P = 0.004$ ($W = 84$) | $P = 0.002$ ($V = 54$) | $P = 0.999$ ($V = 1$) |
| *C. gestroi* | *C. formosanus* | $P = 0.072$ ($W = 70$) | $P = 0.544$ ($W = 49$) | $P = 0.138$ ($V = 39$) | $P = 0.884$ ($V = 16$) |

$T_{L \rightarrow F}$ : transfer entropy from leader to follower, $T_{L \rightarrow F}^S$ : transfer entropy from leader to follower in surrogate datasets. $W$ statistics are provided for Wilcoxon rank-sum test, while $V$ statistics for Wilcoxon signed-rank test.

